# The Role of Sex in the Effects of Smoking and Nicotine in Cardiovascular Function, Atherosclerosis, and Inflammation

**DOI:** 10.1101/2024.02.13.580217

**Authors:** Ann Marie Centner, Abigail Cullen, Leila Khalili, Vladimir Ukhanov, Stephanie Hill, Riley Deitado, Hyun Seok Hwang, Tooyb Azeez, Justin D. La Favor, Orlando Laitano, Michelle S. Parvatiyar, Gloria Salazar

## Abstract

cigarette smoke (CS) invokes an inflammatory response involving increased levels of circulating cytokines and chemokines, vascular dysfunction, and atherosclerosis. The role of sex and nicotine in CS effects in cardiovascular function and atherosclerosis is unexplored. To assess the role of sex, male and female C57Bl/6 WT (wild type) and ApoE^-/-^ mice were exposed to CS and nicotine for 16 weeks to bridge this literature gap. Heart rate and endothelial function were measured in the aorta of WT mice, while plasma levels of lipids, cytokines and chemokines and aortic plaque burden was assessed in ApoE^-/-^ mice. CS increased heart rate to similar levels in both sexes and induced a stronger impairment in endothelial function in males and more plaque in females than nicotine. Females showed a higher necrotic core area at basal compared with males, while males had a higher calcification area than females by CS. Senescence-associated GLB1/μ-galactosidase (SA-GLB1) activity was elevated similarly in both sexes by both treatments. Total cholesterol (TC) was elevated by CS in both sexes. CS increased triglycerides (TG), very-low density lipoprotein (VLDL) and high-density lipoprotein (HDL) only in males and low-density lipoprotein (LDL) only in females. Interleukin 17A (IL17A) was upregulated by CS and nicotine in both sexes, while CS upregulated C-X-C motif chemokine ligand 5 (CXCL5/LIX) and interleukin 1 alpha (IL1A) in males and females, respectively. Additionally, nicotine metabolism showed sex specific responses to nicotine, but not smoking. Overall, we identified sex-specific pro-atherogenic responses to CS in the lipid profile, plaque area and composition and inflammatory markers. Males present a stronger impairment in endothelia dysfunction in WT mice, while females a stronger plaque burden in ApoE^-/-^ mice exposed to CS. Elevated HDL and estrogens in males may offer partial protection against the harmful effects of CS. In contrast, elevated LDL and a pro-inflammatory state may promote a stronger pro-atherogenic phenotype in females exposed to CS.

## Introduction

Atherosclerosis is fueled by modifiable and non-modifiable risk factors, including a Western diet, smoking, genetics, and sex. Cigarette smoking is the number one modifiable risk factor for cardiovascular disease (CVD). Nicotine in cigarettes dysregulates the autonomic nervous system increasing heart rate and blood pressure, which stresses the vasculature and leading to oxidative damage [1]. Nicotine-induced endothelial dysfunction appears to be primarily derived by decreased bioavailability or generation of Nitric oxide (NO).[2] CVD fatality is higher in females, and their risk goes up significantly following menopause, which is partly attributed to a decrease in estrogen levels leading to higher LDL-cholesterol [3]. In addition, women’s cardiovascular system is more sensitive to smoking, as women smokers have more heart attacks than men of a similar background [4]. Human sex differences warrant further studies to elucidate the underlying mechanisms. However, sex-dependent differences in vascular function and atherosclerosis induced by nicotine CS remain to be elucidated as mice studies testing nicotine use mainly males [5–8], while studies testing cigarette exposure are split between sexes [9–12]. The absence of studies using both sexes to assess the impact of nicotine, compared with CS, in CVD has led to the concept that nicotine may be less harmful than CS [9,13]. The purpose of this study was to fill this gap using sex as a variable to assess the effects of CS and nicotine in vascular function and atherosclerosis with the purpose of identifying sex-specific risk factors.

While widely regarded for its contribution to adverse pulmonary function and chronic obstructive pulmonary disorder (COPD), an inflammatory lung disease, smoking also promotes atherosclerosis via inflammatory mechanisms [14]. Damage to the intima is regarded as the instigator of the initial inflammatory response [15]. IL17A is one pro-inflammatory mediator upregulated in our study under both conditions (nicotine and CS). This upregulation is in line with observations from human smokers’ plasma [16]. We noted other changes in inflammation meditators such as CXCL5, which is also upregulated in COPD and atherosclerosis, expressed highly in bronchiolar lavage fluid[17] and plasma [18] from humans. In addition, IL1A – another inflammatory cytokine that is increased during COPD [19] and atherosclerosis [20] and with CS exposure [21]. Lastly, IL1A – regarded as a prominent senescent cell marker [22], its expression may correlate with senescent cell accumulation that occurs during chronic diseases.

During atherosclerosis initiation, vascular smooth muscle cells (VSMCs) undergo a phenotypic switch from contractile to synthetic [23], migrating to the intima where they proliferate. They exhibit reduced markers of differentiation and increased secretion of extracellular matrix (ECM) remodeling factors, including matrix metalloproteinases (MMPs). Specific MMPs, such as MMP2 and MMP9, facilitate VSMC migration [24]. Plaque instability is promoted by MMP secretion, as MMPs degrade ECM components. VSMCs also can differentiate into foam cells, as they engulf oxidized LDL [25] and they can increase plaque stability through secretion of collagen [26]. During advanced atherosclerosis, some VSMCs undergo apoptosis while others become senescent secreting many components that further drive inflammation [27].

As the atherosclerotic plaque develops, fatty deposits increase in size, fibrous and connective tissue is produced, and deposition of calcium occurs [28]. A more robust fibrotic cap and a greater degree of calcification increases plaque stability. On the other hand, unstable plaques are characterized by a thin cap and large necrotic core containing an accumulation of necrotic cells, cellular debris, and lipids [29]. An unstable plaque is generally regarded as more dangerous because of its potential to rupture and travel to other areas of the body and occlude an artery [15]. Sex differences in humans have been noted. Namely, compared to men, women’s developed plaques contain more cellular fibrotic tissue and calcification and fewer non-culprit lesions (e.g., lesions away from the infarct site) [30,31]

Here we show sex-specific differences in endothelial function, plaque, inflammation, and regulators in mice exposed to CS and nicotine. With CS, endothelial dysfunction was more severe in males and plaque accumulation greater in females. CS and nicotine exposure increased necrotic core area and calcification, with male mice experiencing greater calcification. IL17A was the only cytokine increased by CS and nicotine in both sexes. CS also increased LDL and IL1A in females and HDL, CXCL5 and MMP3 in males.

## Results

### Sex-dependent effects of nicotine and cigarette smoke on cardiovascular function

To evaluate the role of sex in the effects of nicotine and smoking, we exposed male and female WT mice to CS and nicotine in drinking water and measured heart rate and endothelial function. Male and female mice exposed to nicotine ate more food in almost all weeks than control and CS groups (Figure S1A and D), while food intake was similar in males and females in the control and CS groups. For water intake (Figure S1B and E), both sexes in the nicotine groups drank less water compared with control and CS-exposed mice. Females tend to drink more water in the smoking group, which reached significance for weeks 4, 5, and 8 compared with control. Significant increases in body weight, compared to week 1, were seen for male controls at week 6, males in nicotine at week 7 and males in smoking at week 13 (Figure S1C). Compared with week 1, males in the control and nicotine groups gained body weigh similarly with ∼20% increases for control and ∼21% for nicotine at week 16. In contrast, smoking only led to a ∼12% increase body weight at week 16 compared with week 1. For females (Figure S1F), changes in body weight were similar for control and smoking groups. Nicotine, however, showed an increase in body weight starting at week 6 compared with week 1, which was significantly higher than the control group from week 13-week 16 (∼26% for nicotine, ∼14% for smoking and ∼10% for control). Although, both sexes ate more food compared with control and nicotine, only females gained more weigh compared with the other two groups.

Serum cotinine in CS-exposed mice was similar between sexes (40.1 ± 18.9 and 54.3 ± 12.3 ng/ml, for males and females, respectively) and was consistent with the levels seen in human smokers (19-50 ng/ml) [32,33] (Figure 1A). Nicotine showed a higer trend in cotinine levels, compared with CS, that was significant only in females. A similar trend was seen when cotinine was adjusted by body weight (Figure 1B).

**Figure 1.**
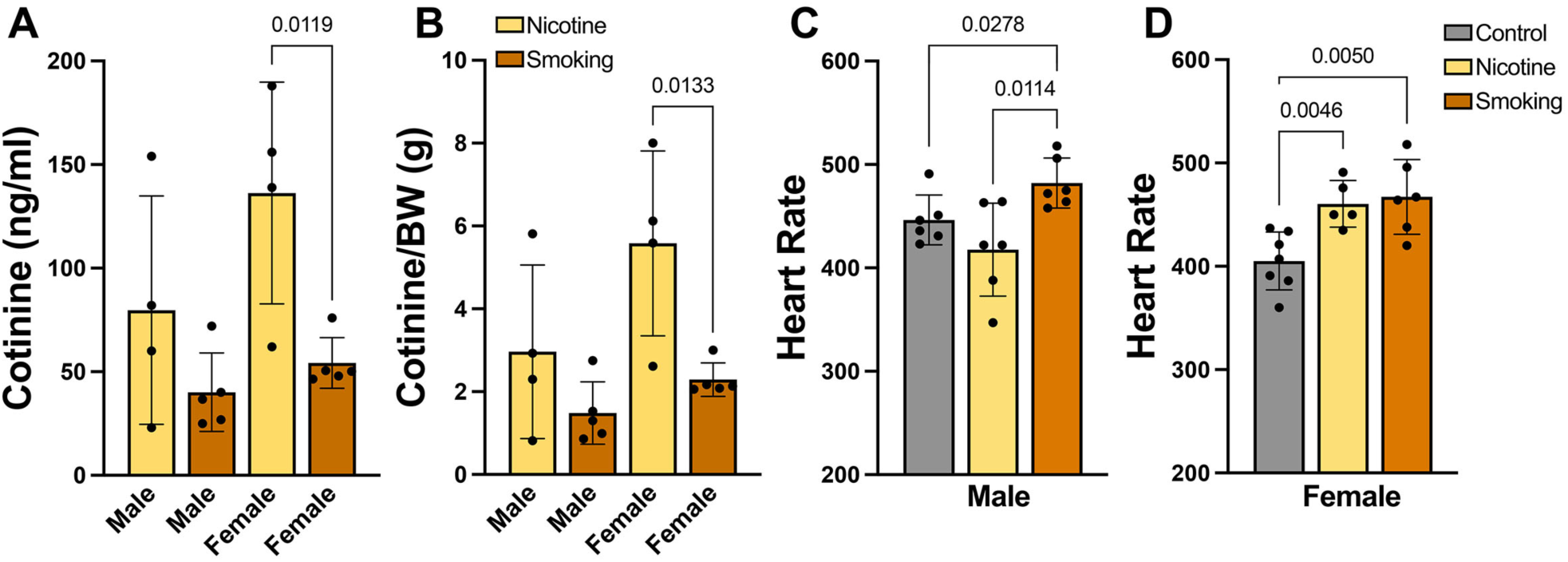
Interaction of sex and treatment in nicotine metabolism and heart rate in C57Bl/6 mice. Mice were exposed to cigarette smoke and 0.2 mg/ml nicotine in drinking water for 16 weeks. Cotinine was measured in serum (**A**) and its levels were adjusted by the mice body weight(**B**). Heart rate was measured for 10 min before sacrifice for males (**C**) and females (**D**).

Resting heart rate was increased in both sexes exposed to CS and only increased in females by nicotine (Figure 1C and D). CS had a greater impact on heart rate in females (∼15%), compared with males (∼8%). To evaluate endothelial function, we induced vasoconstriction with phenylephrine (PE) in aortas and measured endothelium-dependent relaxation to acetylcholine (ACh). Endothelium-dependent relaxation was impaired by CS and nicotine in both sexes (Figure 2A and B). Males however showed a stronger impairment in relaxation by CS than females (Figure 2C). No sex differences were seen for nicotine (Figure 2D). To test whether impairments were endothelium dependent, we induced vasorelaxation with the NO donor sodium nitroprusside (SNP) and observed no major differences among groups (Figs. 2E and F). Modest increases in endothelium-independent relaxation at the lower SNP doses for male CS, female CS, and female nicotine aortas indicates a subtle sensitization to NO, potentially due to a chronic impairment of NO production. There were no group differences in aortic vasoconstriction following 120 mM KCl exposure (Male – Control: 4.41 ± 0.45; Nicotine: 4.23 ± 0.24; CS: 4.96 ± 0.29; Female – Control: 5.11 ± 0.29; Nicotine: 4.38 ± 0.57; CS: 4.58 ± 0.66 mN/mm) or 10 µM PE pre-constriction (Male – Control: 161 ± 8.1; Nicotine: 154 ± 7.2; CS: 172 ± 9.3; Female – Control: 153 ± 13.1; Nicotine: 173 ± 16.9; CS: 151 ± 13.0 % KCl constriction). These data suggest that nicotine mediates the elevation in heart rate and endothelial dysfunction in females, but not in males and that CS has a stronger effect in males, compared with nicotine.

**Figure 2.**
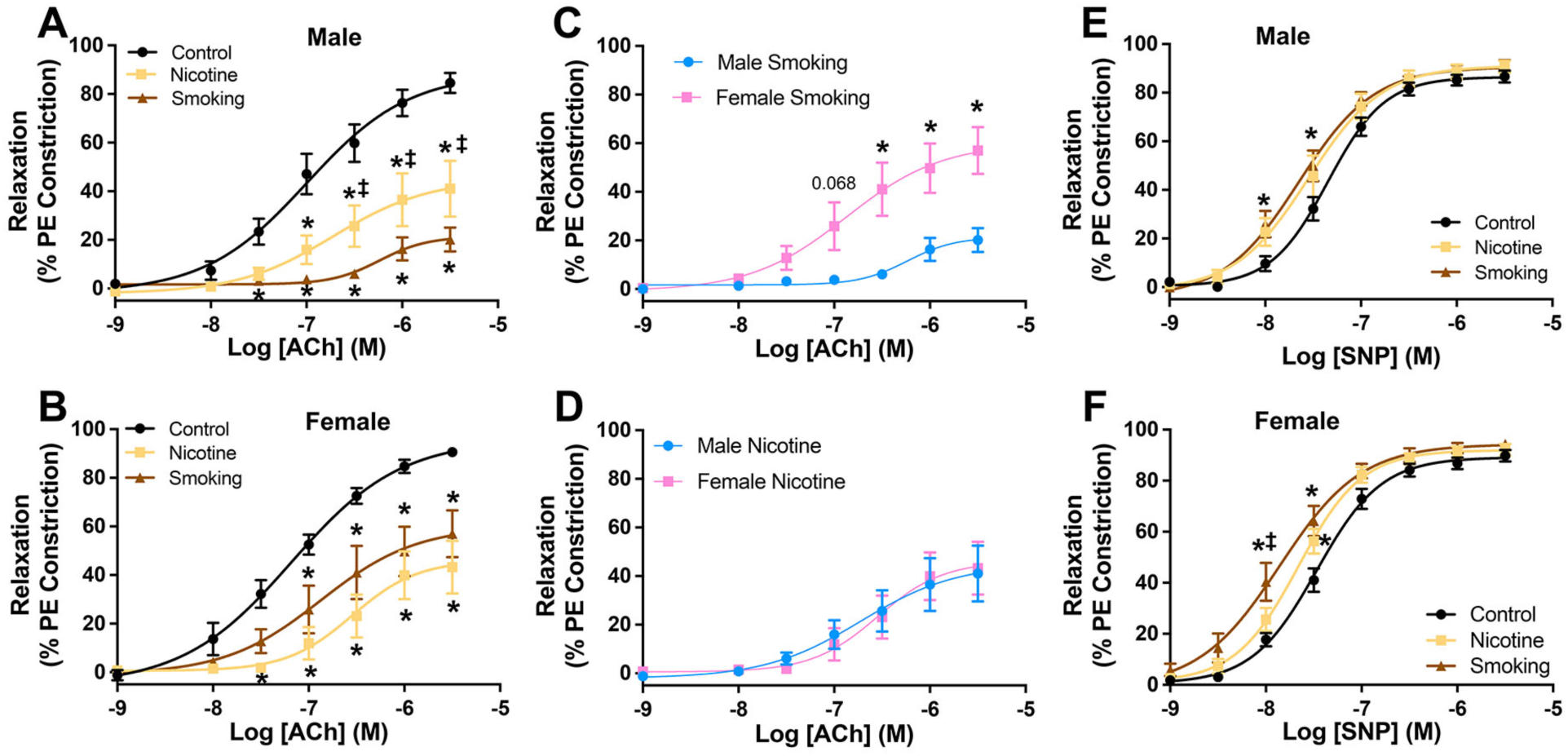
Cardiovascular adaptations to cigarette smoke and nicotine in male and female C57Bl/6 mice. Mice were exposed to cigarette smoke and 0.2 mg/ml nicotine in drinking water for 16 weeks. Endothelial (**A-D**) and smooth muscle function (**E-F**) were assessed in PE pre-constricted aortic rings by cumulative addition of ACh and SNP, respectively. The symbols * and ‡ represent P<0.05, compare with control and between nicotine and smoking, respectively. Data are presented as mean ± SEM.

### Sex-dependent effects of nicotine and CS on atherosclerotic burden

Next, we evaluated sex-dependent differences in atherosclerosis in ApoE^-/-^ mice, a model used extensively to evaluate the effects of smoking in CVD [34]. We exposed male and female ApoE^-/-^ mice to CS and nicotine for 16 weeks and measured body weight, and food and water intake. Male and female mice exposed to nicotine ate more food across all weeks than control and CS groups (Figure S2A and D). While no differences in food intake were seen for males by CS, females showed a tendency toward higher food intake that was significant only in some weeks. For water intake (Figure S2B and E), males in the CS group drank more all weeks. In contrast, females in this group drank more water than the nicotine, but not control. Nicotine groups drank less water only in the first few weeks of treatment, compared with controls. Their water intake remained similar to controls for the rest of the experiment. Compared to week 1, body weight increased in all groups (Figure S2C and F), reaching significance for nicotine after week 7 for males and week 4 for females. Males and females gained more weight for CS and controls after weeks 13 and 14, respectively. For body composition, compared with week 1, males in all groups gained fat, and lost lean mass and total water by week 16 (Figure 3A-C). Females showed no differences in fat or lean mass, but controls had increased total water at week 16 (Figure 3A-C). Free water was elevated in the control group at week 16 in both sexes, which was reduced by both treatments in both sexes (Figure 3D).

**Figure 3.**
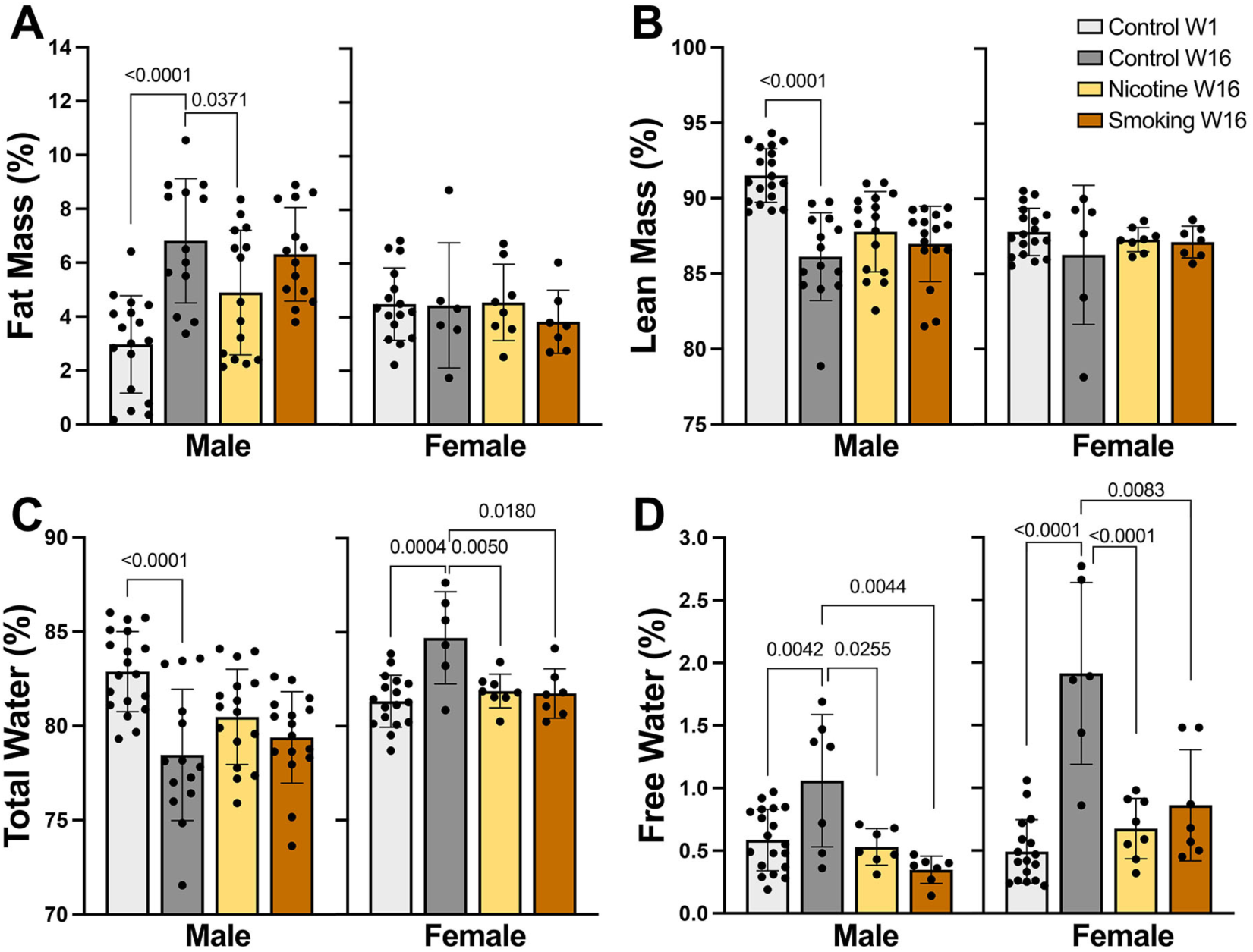
Effect of nicotine and smoking in body composition in male and female ApoE^-/-^ mice. Mice (n=8 per group) were exposed to CS and 0.2 mg/ml nicotine in drinking water or control air for 16 weeks. Fat (**A**), lean (**B**) and total (**C**) and free water (**D**) masses were measured before (week 1) and after treatment (week 16). Data are presented as mean ± SD.

We next compared body composition between sexes (Figure S3). Baseline characteristics (week 1) between sexes differed, as females had more fat mass and lower lean and water masses than males (Figure S3A-C). However, at week 16, females had less fat and more water in the CS group and more water in the control group, compared with males.

Plaque was then assessed by en face analysis in both sexes at week 16 (Figure 4). Compared to control, plaque was higher in the arch of nicotine and CS exposed mice in both sexes (Figure 4A, B, and E), but only females accrued more plaque in the descending aorta by both treatments (Figure 4A, C, and F). The greatest plaque area was seen in the arch (∼39%) and descending aorta (∼6%) of females exposed to CS. Males showed ∼22% in the arch and ∼3% plaque area in the descending aorta with CS. Like plaque content, senescence-associated GLB1/β-galactosidase (SA-GLB1) activity was upregulated in all groups and was higher in females in the CS, compared to the nicotine group (Figure 4D and G).

**Figure 4.**
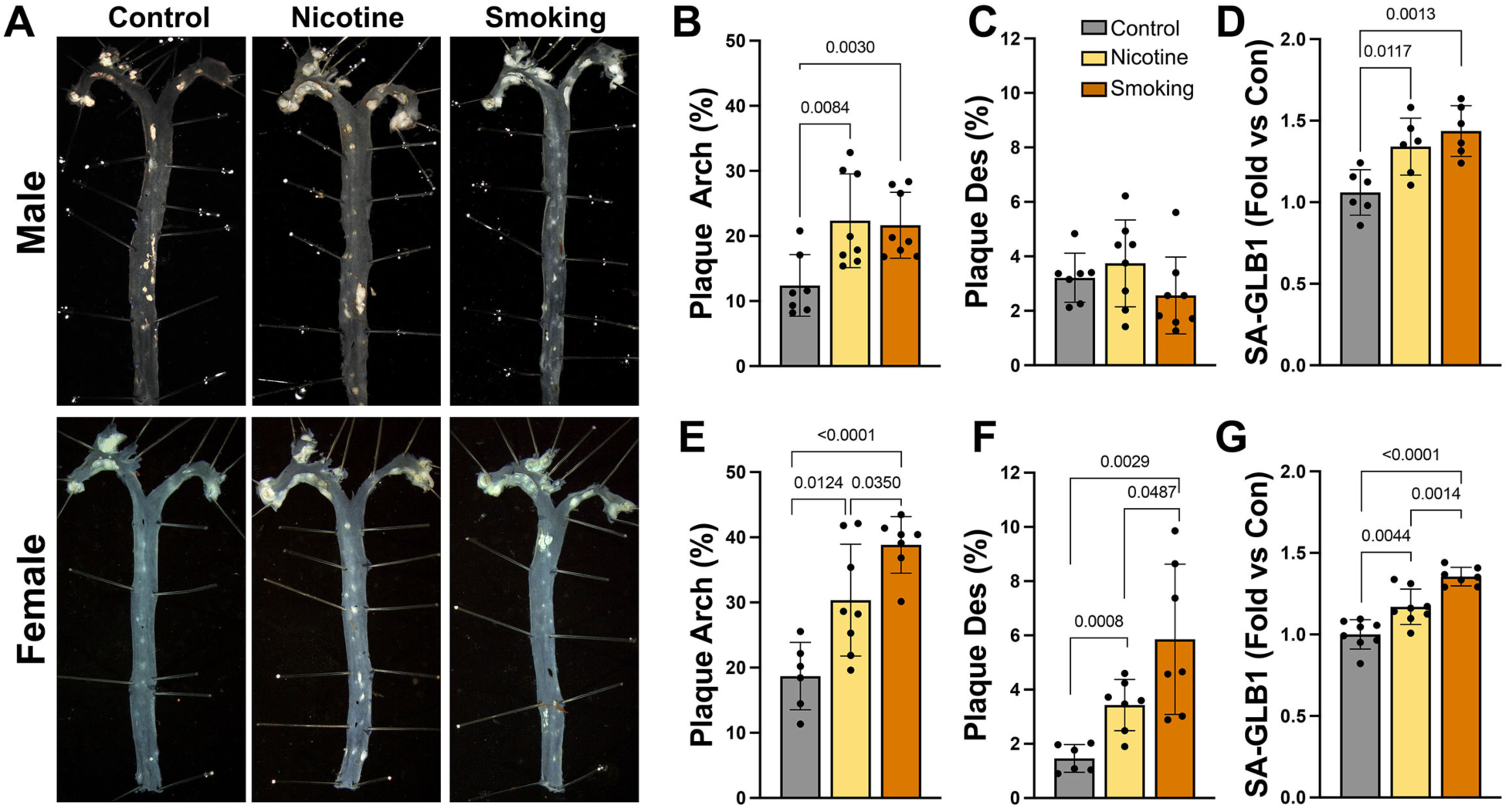
Sex-dependent effects of nicotine and cigarette smoke in plaque in ApoE^-/-^ mice. Male and female mice (8/group) were exposed to 0.2 mg/ml nicotine in drinking water, CS or control water and fresh air for 16 weeks. Aortas were isolated, fixed and opened for plaque quantification using ImageJ (**A**). Plaque was quantified in the arch (**B,E**) and descending (Des) aorta (**C,F**). SA-GLB1 activity was measured using FDG and fluorescence adjusted by the weight of the aorta (**D,G**). Data are presented as mean ± SD.

For plaque composition, both sexes exposed to CS and nicotine experienced similar elevations in necrotic core areas (Figure 5A and B). At basal however, females had a significant higher necrotic core area compared with males. Sex-specific differences were seen for calcification in CS-exposed animals with males showing a higher calcification area compared with females (Figure 5A and C). Calcification was similar in CS- and nicotine-exposed females and in nicotine-exposed males.

**Figure 5.**
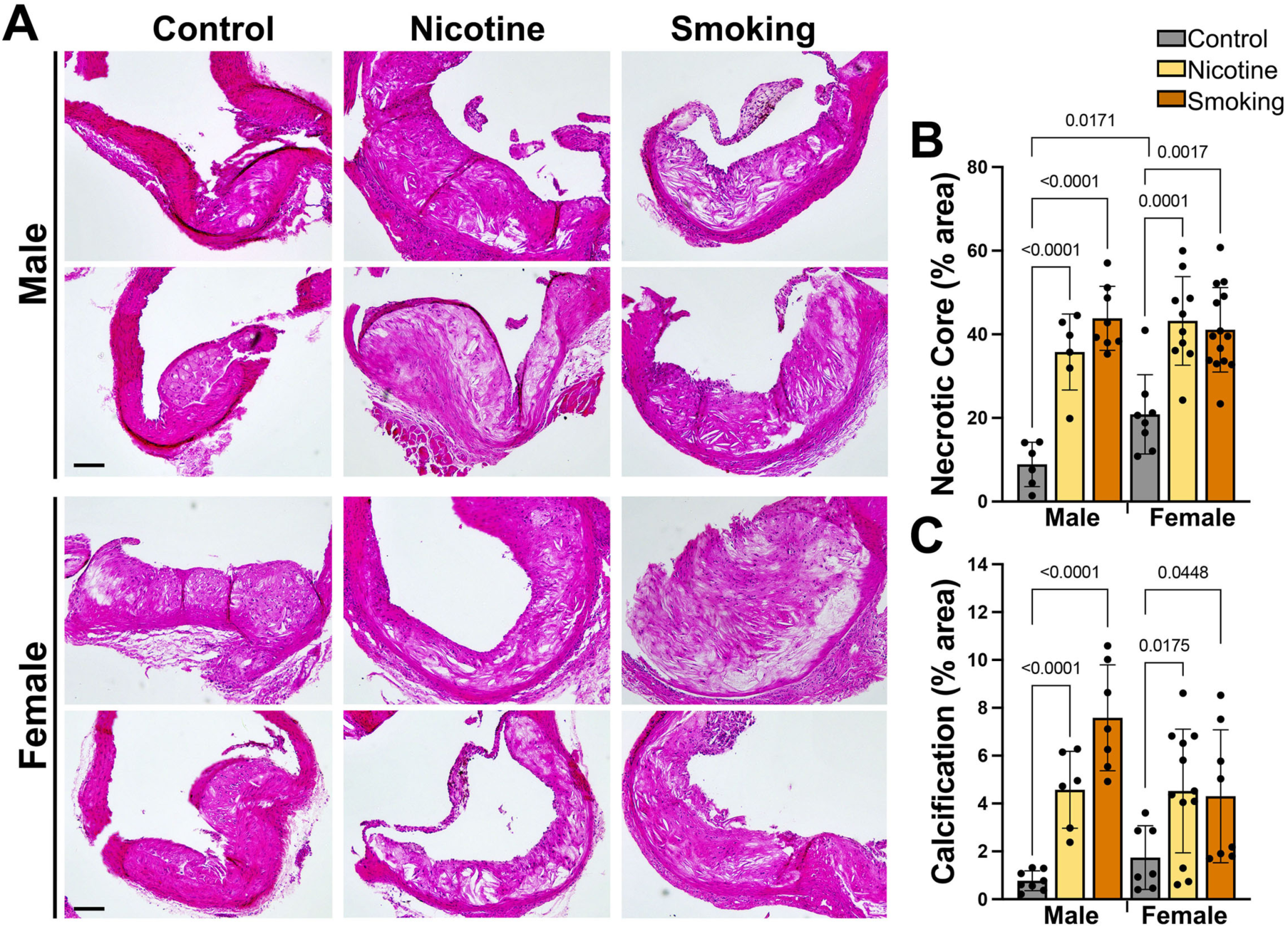
Sex-specific effects of nicotine and cigarette smoke in plaque composition. Male and female mice (8/group) were exposed to 0.2 mg/ml nicotine in drinking water, CS or control water and fresh air for 16 weeks. Aortas were isolated, fixed and stained with H&E (**A**) for measurements of necrotic core (**B**), and clacification areas (**C**) in the arch. Data are presented as mean ± SD. Barr is 100 mm in A.

### Sex-dependent effects of nicotine and CS on circulating lipids, homones and systemic inflammation

Next, we assessed whether plaque accumulation was associated with changes in serum levels of cholesterol, lipoproteins, glucose, liver enzymes, and/or sex hormones. In males, CS increased total cholesterol (TC), triglycerides (TG), HDL, and very low-density lipoprotein (VLDL), compared with control and nicotine (Figure 6A, B, D and E). No changes were seen in LDL, while glucose was increased by CS and nicotine (Figure 6C and F). In females CS increased TC and LDL compared with control and nicotine (Figure. 6A and C). The liver enzymes alanine and aspartate aminotransferases (ALT, AST) were elevated by CS, compared with nicotine in males, but only AST reached significance compared to control in females (Figure 6G and H).

**Figure 6.**
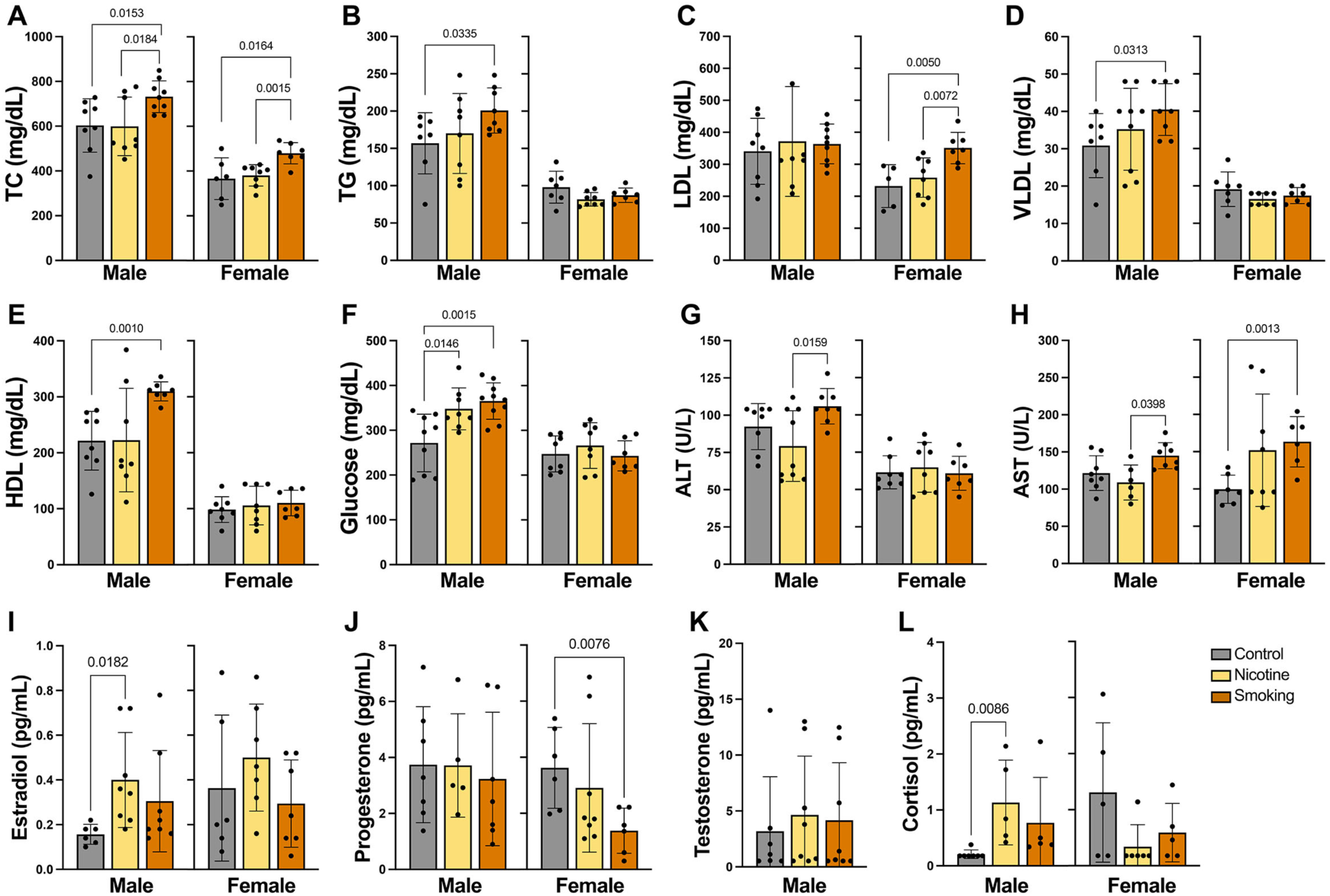
Sex-specific effects of nicotine and CS in lipid and hormone profile in ApoE^-/-^ mice. Male and female apoe^-/-^ (8/group) were exposed to 0.2 mg/ml nicotine in drinking water, CS or control water and fresh air for 16 weeks. Blood serum was used for the analysis of lipids (**A-E**), glucose (**F**), liver enzymes (**G,H**), estradiol (**I**), progesterone (**J**), testeosterone (**K**), and cortisol (**L**). Data are presented as mean ± SD.

Estradiol levels in males were higher in the nicotine and CS groups compared with control, but only nicotine reach significance (Figure 6I). Progesterone and testosterone were not affected by any treatment (Figure 6J and K). In females, estradiol was not different between groups, while progesterone was reduced by CS, but not by nicotine, compared with control (Figure 6I and J). Testeosterone was less than 0.52 pg/mL in all samples and groups in females. Cortisol was elevated in males by CS and nicotine, but only nicotine reached significance. In females, cortisol levels were higher at basal compared with males with no differences between groups (Figure 6L). Thus, reduced estradiol, which protects females from CVD, is likely not the cause of increased plaque by CS or nicotine.

Overall, plaque and lipid profile differed between sexes, as females displayed greater plaque in the control (Figure S3D, arch only) and CS (Figure S3F, arch and descending aorta) groups and a trend towards more plaque by nicotine (Figure S3E, arch). These differences were not associated with more circulating lipids. In fact, females had less TC, TG, HDL, and VLDL than males in all groups and lower glucose in the nicotine and CS groups (Figure S3G-I). Although LDL was elevated by CS in females, its levels were not different between sexes in any groups.

Since the lipid and sex hormone profiles did not completely explain sex disparities nor treatment’s effect on plaque, markers of inflammation (interleukins and CXC and CC motif ligand (CCL) chemokines), CVD markers, and MMPs were examined.

For males, IL17A was the only cytokine upregulated by both treatments (Figure S3H) and nicotine showed a downward trend in CCL2/MCP1 (Figure S3M), compared to control. CS led to higher granulocyte colony-stimulating factor (CSF3), CXCL5 and lower IL-5 and monokine-induced by gamma interferon (MIG/CXCL9), compared to control (Figure S3 A, I, E and J). Similar to males, females showed increased IL17A expression by both treatments (Figure S4H). Bot treatments also reduced IL-5 and IL-9, compared to control (Figures S4E and F). Nicotine upregulated the chemokine keratinocyte chemoattractant/growth-related oncogene (KC/GRO/CXCL1) and CXCL9, while CS increased interleukin 1 alpha (IL1A) and reduced CSF3 and interferon gamma-induced protein 10 (IP10/CXCL10), compared to control (Figure S4L, J, C, A and K).

For CVD markers and MMPs, males showed elevated soluble E-Selectin (SELE) and thrombomodulin (THBD) by CS Figure (S5A and D), while females expressed more SELE in the nicotine group (Figure S5A). Soluble P-Selectin (SELP), soluble intercellular adhesion molecule 1 (sICAM-1) and MMP2 were not affected by treatment or sex (Figure S5B). MMP3 was increased by nicotine in males and females and by CS only in males (Figure S5F). MMP8 was reduced by nicotine in males, and it was not changed by treatment in females (Figure S5G). Pro-MMP9 was not different between groups in males (Figure S5H) and was reduced by nicotine and CS in females reaching significance only for nicotine (Figure S5I).

Next, we compared the expression of these markers between sexes and treatments (Figure 7). Values were calculated as fold change compared with males in the control group. A robust upregulation of IL1A, IL17A and CXCL5 was seen in females, compared with males in all groups (Figure 7A-C). At basal females also had elevated CSF3, IL4, IL5, IL13, CXCL10, and CXCL10 and reduced CCL11 and IL-9, compared to males (Figure 7D). Similar trends were seen for nicotine except for IL5, which was similar between sexes and CCL2, which was upregulated in females compared with males (Figure 7E). Different from control, smoking led to similar levels in CSF3, IL4 and IL5 (Figure 7F).

**Figure 7.**
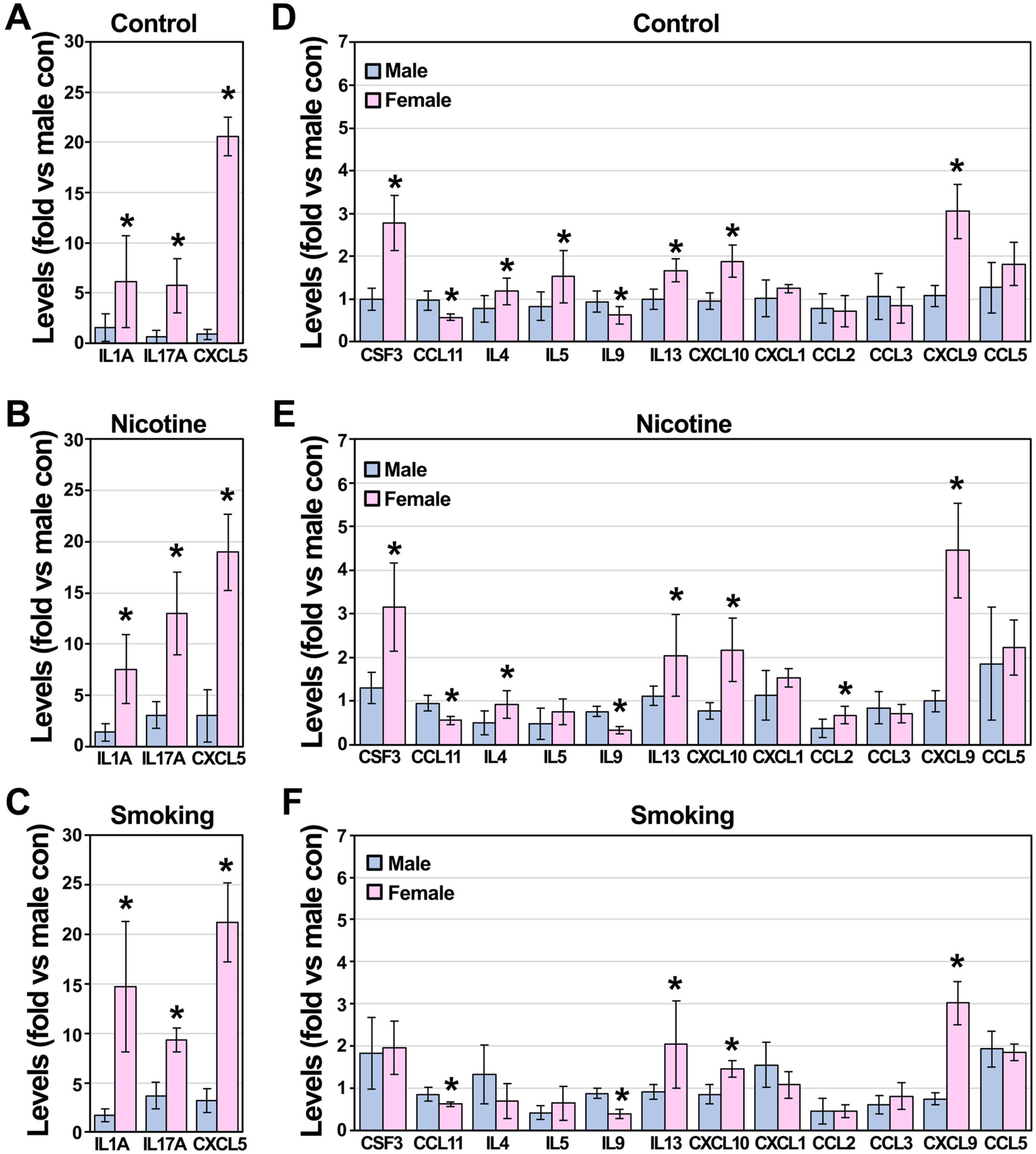
Sex differences in the expression of inflammatory markers. Cytokines and chemokines were analyzed in plasma from males and females controls (**A,D**) and exposed to nicotine (**B,E**) or CS (**C,F**) using a Luminex system. Expression was calculated as fold change compared with male control. Values are expressed as mean ± SD.

Regarding CVD makers and MMPs, females had lower SELP and THBD compared with males in all groups (Figure 8A-C). Although, SELE level was the same between sexes at baseline, its level was higher in females exposed to nicotine and in males exposed to CS (Figs 8A-C). No differences between sexes were seen in MMP2, 3 and 8 levels, except for MMP8, which was reduced by CS in females (Figure 8D and E). Only MMP8 was reduced in females by CS (Figure 8F). In contrast, pro-MMP9 expression was about 40-fold higher in females at basal (Figure 8D) and about 12- and 5-fold higher in the nicotine (Figure 8E) and CS (Figure 8F) groups, respectively than males.

**Figure 8.**
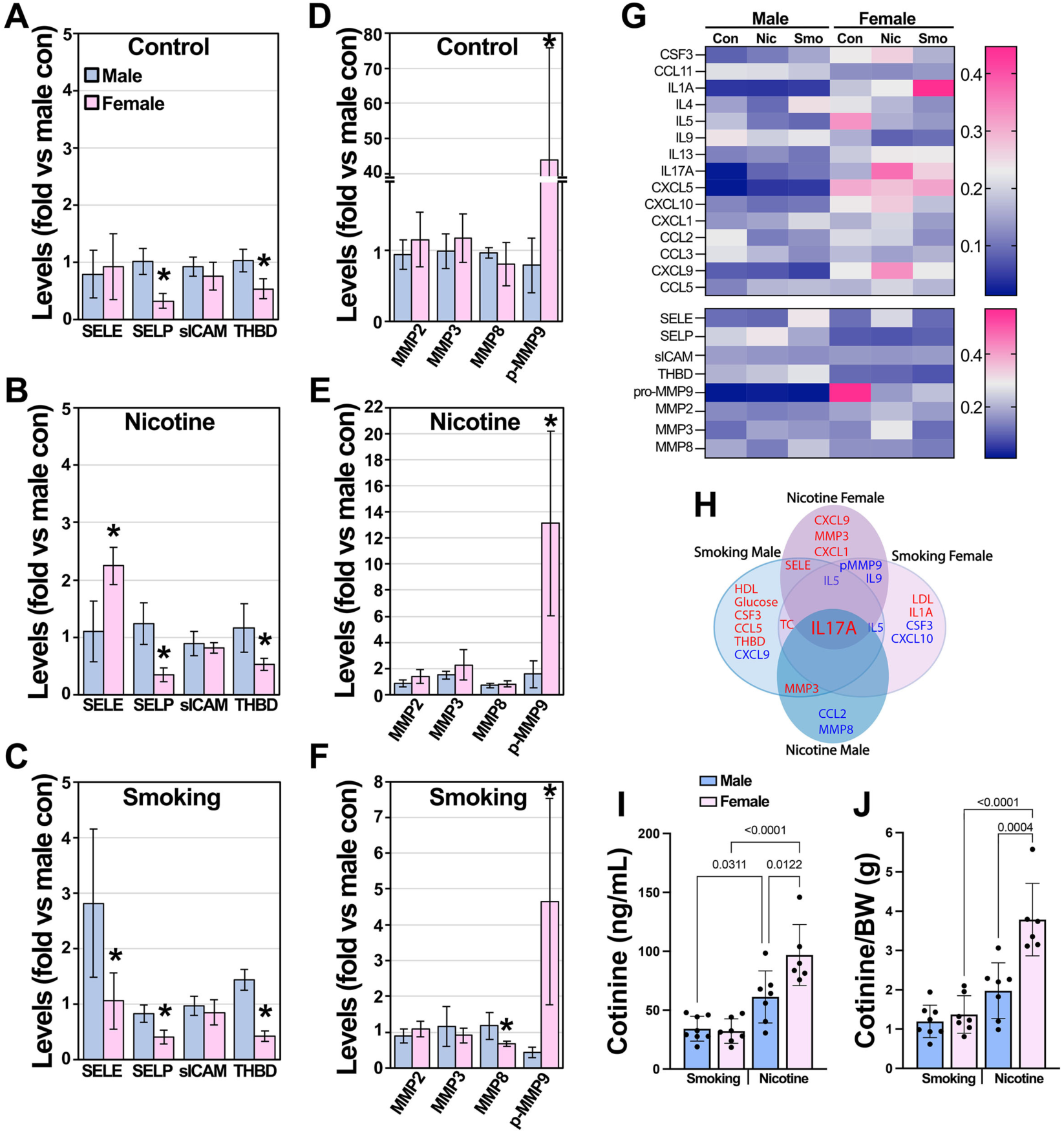
Sex differences in the expression of markers of inflammation, CVD, and ECM remodeling. CVD markers (**A-C**), and MMPs (**D-F**)were analyzed in plasma from male and female controls and those exposed to CS and nicotine using a Luminex system. Expression was normalized in the heat map (**G**). Illustration of the major markers upregulated (red) and downregulated (blue) in males (light blue) and females (pink) by treatment (**H**). Cotinine was measured in plasma (**I**) and was adjusted by body weight (**J**). Values are expressed as mean ± SD.

The heat map in Figure 8G shows that the most prominent changes are seen for IL1A by smoking in females and pro-MMP9 expression in control females. Altogether, females present a pro-inflammatory atherosclerotic state with increased IL1A, IL17A, and CXCL5, higher LDL levels and plaque (Figure 8H). IL17A was the only cytokine upregulated by both treatments and in both sexes. Differential nicotine levels/metabolism explain, in part, sex-differences in plaque since cotinine, a nicotine metabolite, was higher in the nicotine groups with females showing a stronger upregulation compared with males; however, cotinine was similar in the smoking groups (Figure I). After adjusting the cotinine levels by the weight of mice no significant differences were seen between males in the smoking and nicotine groups, but cotinine remained elevated by nicotine in females (Figure 8J)

Next, we investigated potential correlations between plaque (arch and descending aorta (DA)) and makers of inflammation, and CVD, MMPs, lipid profile, glucose, and hormone levels for all animals and treatments combined (**Table 1**). For plaque in the arch, a strong positive correlation was seen for IL1A and plaque in both sexes. A moderate negative correlation close to significance was seen for IL5 and CXCL10 in males. For females, strong negative correlations were seen for CSF3, IL5, IL9 and MCP1 while a moderate negative correlation was seen for TG. Several very strong (p<0.001) positive correlations in sex effects were identified, including TC, LDL, HDL, CSF3, IL5, MCP1, and SELE followed by strong positive correlations in VLDL, glucose, IL4, CXCL1, CXCL2, CXCL9, CXCL10, CCL3, CCL5, SELP, sICAM, pro-MMP9, cortisol, estradiol, progesterone, MMP2 and pro-MMP9, and moderate positive correlations in TG, IL13, CCL11 and THBD. For plaque in the descending aorta, males showed no significant association between plaque and any marker tested. In females, IL1A showed a moderate positive correlation and MPC1 a moderate negative correlation with plaque. A trend towards a positive correlation was seen for LDL and IL13 and a trend towards a negative correlation for IL5. No sex effects were identified in the descending aorta.

**Table 1.**
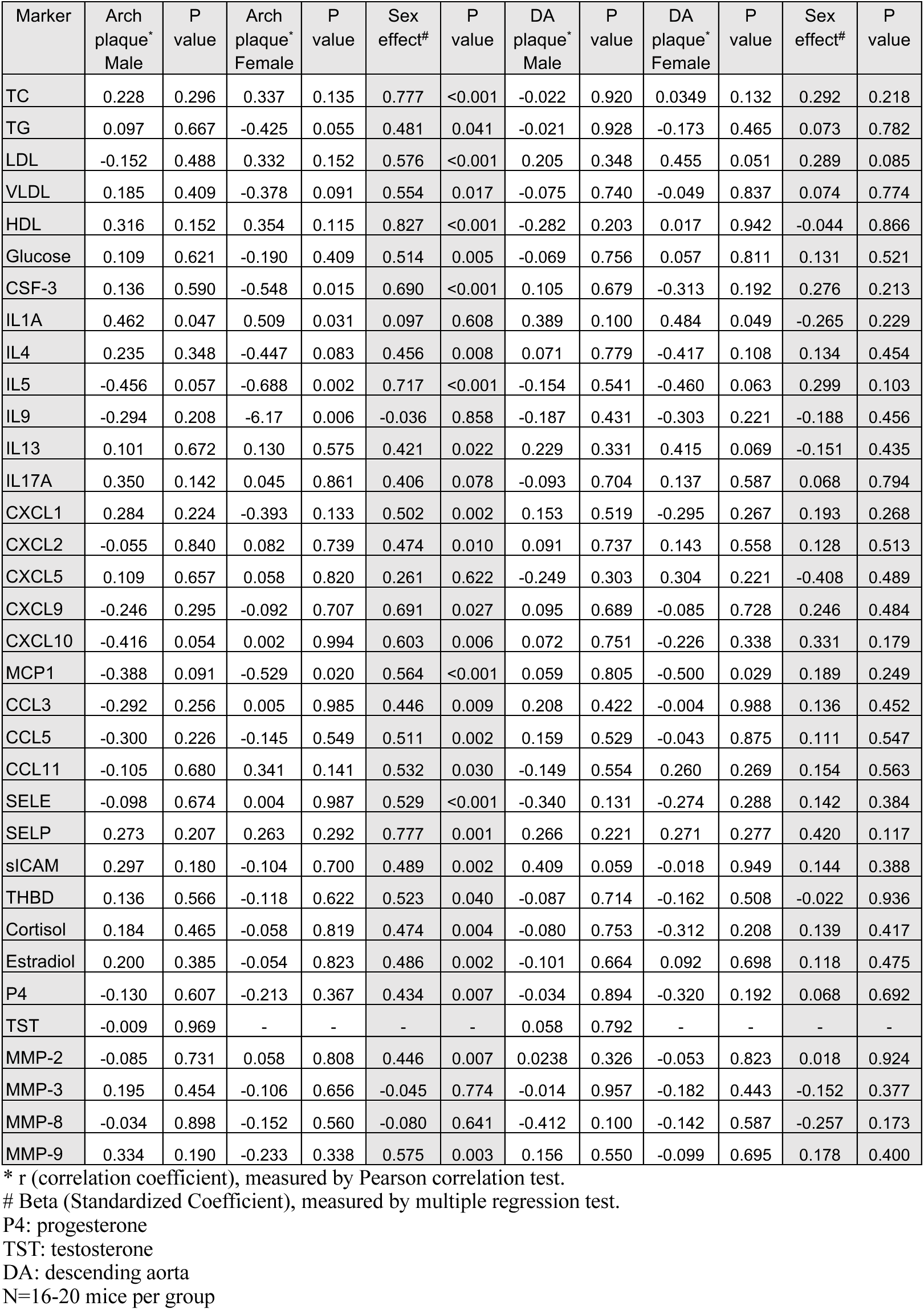
Correlation analysis for plaque and lipids, inflammatory and CVD markers, MMPs and hormones.

## 3. Discussion

Smoking is known to increase atherosclerosis, but whether one sex is more prone to nicotine or CS’s effects on the cardiovascular system is yet to be determined. There is a current gap in the literature, as studies assessing plaque and other cardiovascular outcomes through exposure to nicotine in drinking water or CS lack concurrent utilization of both male and female WT and ApoE^-/-^ mice. Female ApoE^-/-^ mice are the preferred sex in many smoke-related studies because they are more sensitive to CS. Here we used male and female C57Bl/6 and ApoE^-/-^ mice and report sex-dependent effects of CS and nicotine in drinking water in heart rate, endothelial dysfunction, body composition, plaque burden and inflammation in vivo.

In male WT mice, CS had the strongest effect in resting heart rate and endothelial function, while in females both treatments resulted in similar responses. This suggests that nicotine mediates the elevation in heart rate and endothelial dysfunction in females [35]. In addition, CS appears more harmful than nicotine in males while both treatments are equally harmful in females. Differences between the male and female response to smoking may be attributed to the link between sympathetic nervous system activation (SNA) and the ovarian cycle in females that peaks during the high hormone level mid-luteal phase. This fluctuation in muscle SNA was found in never-smokers, but not present in women who smoke [36]. This finding suggests that the increased resting heart rate in female smoked mice could be due to an alteration of normal SNA alterations associated with the ovarian cycle. Nicotine in drinking water (100 mg/L) has been used in mice [37] and rats [38] to show impaired endothelial function. These studies were done only in males and with high fat diet (HFD). With respect to CS, two studies using male WT mice showed altered endothelial function after 32, but not 16 weeks, of whole-body mainstream CS exposure. One study used the same smoking system we used (SCIREQ InExpose) [39] and the other [40] a TE-10 (Teague Enterprises) system. Females were not used in these studies. Although the conclusion of these studies is similar to ours, we observed a robust alteration in endothelial function by CS at 16 weeks of treatment.

Differences in plaque between sexes are observed in animal models of this disease, including ApoE^-/-^ mice in chow diet and HFD [41]. Many studies showed more plaque in females [42–45] and few saw more plaque in males [46]. In our study, only females exposed to CS accumulated more plaque in the arch, compared with nicotine, and in the descending aorta with both treatments. Thus, females may be at risk of more severe CV events such as stroke as descending aortic plaque accumulation is closely correlated with incidence of a CV event [47]. In addition to changes in plaque localization, sex-differences were seen in plaque composition. Namely, while exposure to CS and nicotine increased necrotic core and calcification, the latter was much more pronounced in male mice, indicating enhanced plaque stability. This is in contrast with a previous study in which ApoE^-/-^ male mice were fed a high cholesterol diet and exposed to CS (5 days per week for 8 weeks). In this study, plaque had higher levels of tissue factor, indicative of thrombus formation and a plaque more vulnerable to rupture [48].

Interestingly, in our study the increase in plaque by nicotine exposure was not correlated with changes in lipid profile, suggesting that other components in cigarettes may regulate lipoprotein metabolism. In fact, exposure to acrolein, an aldehyde in CS, is enough to increase circulating cholesterol and TG in ApoE^-/-^ male mice [49]. LDL/HDL levels are relevant to CVD since in humans the majority of cholesterol is carried by LDL lipoproteins [50], which increases the risk of developing atherosclerosis. In contrast, in rodents most of the circulating cholesterol is carried in HDL [51] making rodents resistant to this disease. In our study, CS increased HDL only in males and LDL only in females. Thus, HDL could be part of a protective mechanism to prevent excessive plaque accumulation in males, while increased LDL could explain, in part, the higher plaque burden in females exposed to CS. Additionally, estradiol was upregulated by CS in male mice, which is in agremment with observations in male smokers [52].

Fueled by similar risk factors, diabetes, obesity, and CVD are often comorbidities [53]. Only males exposed to CS and nicotine had higher glucose, suggesting that impaired glucose metabolism may contribute to their plaque development. Females had more fat and less lean and water masses at the beginning of the study. At 16 weeks, males in all groups gained more fat and lost lean mass and water reaching similar levels to females. Overall, body composition changes do not appear to mediate sex-dependent differences in plaque by treatment.

Nicotine reduces food intake and body weight gain in mice [54]. In contrast, we observed an increase in body weight and higher food intake in both sexes exposed to nicotine. The differences in the study outcomes could be due to study duration, the route of administration, and nicotine concentration. In the study by Mineur et al [54] nicotine was injected via IP once a day per 30 days and only males C57Bl/6 mice were used. In ApoE^-/-^ mice, Gao et al [55] exposed mice to nicotine via osmotic minipumps for 12 weeks and observed an inhibition of body weight gain induced by HFD. Animal sex for this study was not disclosed. We also observed higher water intake in males, but not in females exposed to CS. The cause of this behavior is unknown. Mice on CS could have experienced nose and throat dryness and drinking more water might alleviate these symptoms.

IL17A emerged as the only inflammatory cytokine increased with both treatments in both sexes. A weak correlation with plaque was seen for males and females in the arch and for females in the descending aorta. IL17A is elevated in plasma of human smokers [56] and in human plaques [57]. However, in animal models the literature is mixed in showing that IL17A exerts atheroprotective [58,59] and pro-atherogenic [60,61] effects. There is also strong evidence of IL17A involvement in other diseases such as psoriasis, arthritis, asthma, and inflammatory bowel disease [62]. Th17 helper cells secrete CSF3 [63] and IL17A, the latter stimulating CXCL5 expression [64]. CXCL5 is a chemokine that attracts neutrophils and promotes inflammation through the phosphoinositide-3-kinase (PI3K)/nuclear factor kappa-light-chain-enhancer of activated B cells (NF-κB) pathway. This mechanism may explain the concomitant upregulation of CSF3, CXCL5 and IL17A in males. However, it is unknown why IL17A, but not CSF3, and CXCL5, was upregulated in females.

IL1A is required for dendritic cell recruitment and activation [65], which is associated with COPD disease severity [66] and is also present on the surface of senescent cells as part of their secretory phenotype [67]. IL1A is linked to inflammation in atherosclerosis [20]; produced primarily by foam cells in the growing plaque and acting as a regulatory protein in disease progression [68]. Higher IL1A in females may indicate a more advanced lesion in which senescent VSMCs release senescence-associated secretory phenotype (SASP) factors that further inflammation. In contrast, in males, higher CCL5, SELE, and THBD with CS may indicate more platelet aggregation and higher monocyte differentiation and macrophage infiltration in growing lesions.

Interestingly, nicotine upregulated more inflammatory markers than CS in females, including CXCL1, CXCL9, and SELE. CXCL1 is produced by Th17 cells and acts to further leukocyte infiltration [69]. CXCL9, produced by IFN-γ stimulation, acts as a chemoattractant for active T cells, while sE-selectin is an adhesion molecule activated by thrombosis. This may indicate advanced monocyte, macrophage, and endothelial cell activation [70]. Nicotine however has varied effects on neural circuits, which can alter immune cell function by interacting with their receptors [71]. The influence of nicotine on female immune cell function warrants further study and may contribute to the sex differences seen in this study.

MMP3 upregulation with nicotine (both sexes) and CS (males) may indicate increased ECM degradation and a more unstable plaque phenotype as MMP3 can activate MMP9 [72]. MMP3 also facilitates perclecan breakdown in ECM basement membranes releasing fibroblast growth factor (FGF). MMP2 and -9 degrade decorin to release transforming growth factor (TGF)-1, which is further activated by these MMPs into TGF-β1 [73]. The decrease in the inactive pro-MMP9 in females may indicate higher active MMP9 and a more unstable plaque phenotype. However, the level of MMPs in blood may not reflect differences in plaque stability. While no differences were seen in necrotic core areas between treatments and sexes, calcification was higher in males in the CS group. Analysis of the expression of MMPs and inflammatory markers in the aorta are needed to further delineate the role of these markers in CS and nicotine effects.

Altogether, most of the factors tested showed a higher correlation with plaque in the aorta in females compared with males (TC, LDL, HDL, CSF3, IL5 and others) in ApoE^-/-^ mice. In the absence of an altered lipid profile in WT mice, males showed a stronger endothelial dysfunction towards CS than females, suggesting that sex, and cardiometabolic imbalances contribute to vascular dysfunction and plaque accumulation. Thus, both sexes must be examined in parallel for each individual cardiovascular outcome to assess the severity of smoking over nicotine *in vivo*. Future studies should test the effects of nicotine and CS *in vitro* in aortic endothelial and VSMCs to further elucidate sex differences and the mechanisms by which CS promotes atherosclerosis.

Understanding the sex differences in disease pathogenesis is important for translating laboratory discoveries into clinical real-life settings. Smoking has long been regarded as an exacerbator of atherosclerosis and reduces health- and lifespan [74]. However, many smokers, especially those with chronic diseases exacerbated by smoking, find it difficult to quit [74]. While their addiction to nicotine is strong, they may be willing to make other lifestyle changes. Findings from our study indicate higher HDL in males compared to females may confer protection against atherosclerosis. In humans, females on average have higher HDL, a separation that begins and persists through adulthood and old age [75]. Targeting cholesterol values, markers which many people receive testing for at yearly routine check-ups, is an actionable intervention. One way to increase HDL is through dietary changes, which could delay atherosclerosis progression. Thus, male smokers could be advised to increase omega-3 fatty acids through consumption of fatty fish or a fish oil supplement [76]. Similarly, LDL reduction could be another strategy to decrease atherosclerosis risk. LDL increases in both sexes throughout the lifespan. LDL is lower in females in adulthood, but post-menopausal women have heightened LDL compared to males [75]. Reduction of saturated fats could be recommended for patients with heightened LDL. Altogether understanding the interplay between internal and external risk factors – not only lipids but inflammation markers and body composition in tandem with smoking and nicotine use is important to inform patient recommendations, reduce disease risk and progression, and improve health-span and lifespan. This study also highlights the consequences of nicotine-replacement therapy, used to quit smoking. We show that nicotine alone promotes atherosclerosis by mechanisms that seems to differ from smoking since nicotine showed no effect on the lipid profile but lead to a more robust pro-inflammatory response specially in females. Thus, targeting the lipid profile maybe more beneficial for smokers than vapers using nicotine or patient using nicotine replacement therapy in the form of nicotine patches or oral administration (gums, lozenges). Future studies are needed to evaluate the interaction of sex and vaping in atherosclerosis.

## Materials and Methods

### Animal models

Animal experiments using ApoE^-/-^ and C57BL/6 (WT) mice from Jackson Laboratory mice were performed in compliance with the Institutional Animal Care and Use Committee of Florida State University. Animals were euthanized using CO2. Mice were singly housed to record food and water consumption as well as body weight. Body weight was assessed twice weekly, while food and water were measured three times per week. C57Bl/6 WT were 10 weeks old, and ApoE^-/-^ were 13-16 weeks old at the beginning of experiments. Each experimental group had at least 8 mice. Animals were randomized into groups, but groups were not blinded. Male and female WT and ApoE^-/-^ mice were exposed to either cigarette smoke (7 research cigarettes, 1R6F 0.721 mg nicotine) 5 days per week or nicotine in drinking water (0.2 mg/ml) 7 days per week for 16 weeks. The 1R6F reference cigarettes were obtained from the Center for Tobacco Reference Products at the University of Kentucky. See cigarettes characteristics in Table S1.

### InExpose exposure system

The InExpose system (SCIREQ) is a computerized smoke/aerosol/air exposure system featuring a carousel with a capacity of up to 16 mice (25-80g each). Mice received cigarette smoke via this system, and were housed individually so body weight, food and water could be monitored and recorded. Animals were trained for 5 days prior to the start of the experiment by placement in the system for the same duration as smoke exposure. They received constant fresh air for 48 min. Animals were placed in the system and exposed to whole body aerosol for 48 min/day. 1R6F cigarettes were used, with smoke delivered in a 35-ml puff of 2 s duration followed by 58 s of fresh air. The treatment was 5 days a week for 16 weeks. The cigarettes were placed in a cigarette smoking robot (CSR), with a holding capacity of 24 cigarettes. An automatic lighter ignited each cigarette, and smoke was delivered through a series of tubing and pumps to the animals.

### Nicotine in drinking water

Mice received 0.2 mg/mL nicotine in drinking water as reported [77] every day for 4 months. Nicotine was prepared in acidified water, (pH 4.0) and bottles were weighed at least 3 times per week and were changed biweekly with freshly prepared nicotine-containing water.

### Resting heart rate measurement in anesthetized mice

At the end of 16-week treatment, resting heart rate (HR) was measured as previously described [78]. Briefly, mice were anesthetized with 2% isoflurane with 100% O2. Resting electrocardiogram (ECG) was recorded and calculated for 10 min using two channel electrodes probes (PowerLab, ADInstruments, Colorado Springs CO, US).

### Cotinine and other plasma measurements

Blood was collected from the left ventricle after animals were sacrificed using CO2. Blood was allowed to coagulate at room temperature and collecting tubes centrifuge for 5 min at 5,000 x g. Serum was collected and stored at -80°C. Plasma cotinine, a long-lasting primary nicotine metabolite, was assessed from serum samples taken from mice exposed to nicotine in drinking water and CS. A commercial Cotinine ELISA kit (Origene) suitable for mice was used to quantify blood levels, requiring 10 µl serum from each animal.

A lipid plus kit designed to work in an Abaxis Piccolo XPRESS chemistry analyzer was used to quantify the following metabolites: total cholesterol (TC), HDL, triglycerides (TG), ALT, AST, GLU, LDL, and VLDL. Samples were diluted 1:3 to 1:4 for females and 1:6 to 1:8 for males in phosphate-buffered saline (PBS).

### Measurements of endothelial function

Aortas were dissected under a dissection microscope and cleaned in chilled Krebs solution of the following composition (in mM): NaCl 130, KCl 4.7, KH2PO4 1.18, MgSO4, 1.18 NaHCO3 14.9, D-glucose 5.6, CaCl2 1.56, and EDTA 0.03 in distilled water. A 2 mm ring of the atherosclerosis-prone segment of the proximal thoracic aorta [79] was cut and mounted on 200 µm pins in a DMT 620M myograph system (Danish Myo Technology, Aarhus, Denmark) for isometric tension measurement. Rings were bathed in Krebs solution maintained at 37°C and aerated with bubbled a 95% O2 and 5% CO2 mixture. Rings equilibrated for 1 h, then stretched to a resting tension of 10 mN, followed by an additional 1 h of equilibration. Tissue viability and contractile function were tested with high potassium (120 mM) Krebs solution, with KCl substituted for NaCl. Following successive washes with Krebs solution to reestablish a stable resting tension, rings were constricted with 10 µM phenylephrine (PE), and endothelium-dependent relaxation was tested with a cumulative dose-response (0.001-3.0 µM) to acetylcholine (ACh). Following successive washes with Krebs solution, rings were constricted with 10 µM PE, and endothelium-independent relaxation was tested with a cumulative dose-response (0.001-3 µM) sodium nitroprusside (SNP). Data were continuously recorded with LabChart v8 software (ADInstruments). Relaxations to ACh and SNP were normalized as a percentage restoration to the resting tension from the PE pre-constricted value [80].

### Aortic plaque assessment

Aortas of experimental mice were isolated, cleaned of periadventitial fat, fixed on 0.2% glutaraldehyde in PBS and opened longitudinally for plaque analysis, as reported [81]. Next, aortas were pinned to black wax in a petri dish before photograph and quantification of plaque. Quantification was done by determining the plaque area, compared to the total aortic area using ImageJ software. Briefly, each aorta was traced with the freehand selection of the software. After exclusion of the outside area of the picture, each image was adjusted using the threshold icon until all plaque is white and the rest of the tissue is black. Then, images were inverted showing plaques in black, allowing for quantification with the “analyze particle” feature of the software. Percent plaque was calculated for the arch and descending aorta.

### SA-GLB1 assay

Cellular senescence was determined by quantification of SA-GLB1 activity in mouse aortas using the X-gal analogue FDG, as previously described [82]. Aortas were fixed in 0.2% glutaraldehyde in PBS overnight, then washed in PBS and incubated phosphate buffer (4.8 mM Na2HPO4, 35.2 mM NaH2PO4, pH 6.0) containing 150 mM NaCl, 2 mM MgCl2, 5 mM K3Fe(CN)6, mM K4Fe(CN)6, and 0.05 mM FDG (Ex/Em: 485/515) for 12 h at 37°C. Fluorescence intensity was measured in a Synergy H1 multi-mode plate reader (BioTek Instruments) and was adjusted according to the aorta weight.

### Magnetic bead assay

Magnetic bead kits including Mouse Cytokine/Chemokine, Multispecies Hormone Panel, Mouse MMP Panel, and Mouse CVD Panel were obtained from Sigma-Aldrich to assess relevant blood markers. Plasma samples were diluted (1:1 with provided Assay Buffer) and 96-well plates processed as according to manufacturer’s instruction. Plates were read using a MAGPIX system (Luminex Corporation).

### Quantification and statistical analysis

For animal studies, 8 mice were used per group. Student’s t-test was used for 2 group comparison (males vs females). Group differences for 3 groups including vascular reactivity were determined by two-way repeated measures ANOVA followed by Tukey’s multiple comparisons post hoc analysis if main effects were detected with GraphPad Prism v9 software. Values were reported as mean ± standard deviation of mean (SD) or standard error (SE). *P* < 0.05 was considered statistically significant. Normality in data distribution was determined by skewness, measured using the Shapiro-Wilk score and kurtosis using the Kolmogorov-Smirnov score, which indicate normally distributed data. Data analyses were performed with the IBM (International Business Machines) SPSS 25 (Statistical Package for the Social Sciences version 25) computer program. Values 2-fold from the mean were considered outliers and were removed from data analysis. For correlation analysis, prior to analysis, data underwent screening for missing values, outliers, and adherence to normality assumptions. Bivariate relationships were explored using Pearson correlation coefficients, examining the interplay between lipid profile, glucose, inflammatory markers, cardiometabolic markers, and hormones level and plaque size for different genders. Furthermore, a multiple regression analysis was executed to discern the combined impacts of each biomarker level and gender on plaque size, with gender treated as a categorical variable (male=0, female=1). All statistical analyses were performed using IBM SPSS Statistics (version 29, IBM Corp., Armonk, NY), with a significance level set at α = 0.05.

## Author Contributions

For research articles with several authors, a short paragraph specifying their individual contributions must be provided. The following statements should be used “Conceptualization, G.S., A.M.C., and A.E.C.; methodology, G.S., A.M.C., A.E.C., V.U., L.K, S.H., R.D., H.S.H., T. A., J.D.F., O.L., M.S.P.; software, G.S. and A.M.C.; validation, G.S. and A.M.C.; formal analysis, G.S., H.S.H, L.K., J.D.F and A.M.C.; investigation, G.S. A.M.C., A.E.C., V.U., L.K., T.A..; resources, G.S.; writing—original draft preparation, G.S. and A.M.C.; writing— review and editing, G.S., A.M.C. J.D.F and M.S.P; visualization, G.S. and A.M.C.; supervision, G.S.; project administration, G.S.; funding acquisition, G.S. All authors have read and agreed to the published version of the manuscript.

